# Sting orchestrates the crosstalk between polyunsaturated fatty acids metabolism and inflammatory responses

**DOI:** 10.1101/2020.12.22.423950

**Authors:** Isabelle K. Vila, Hanane Chamma, Alizée Steer, Clara Taffoni, Line S. Reinert, Evgenia Turtoi, Mathilde Saccas, Johanna Marines, Lei Jin, Xavier Bonnefont, Soren R. Paludan, Dimitrios Vlachakis, Andrei Turtoi, Nadine Laguette

## Abstract

Inflammatory disorders are major health issues in which immune function and metabolic homeostasis are concertedly altered. Yet, the molecular mechanisms coordinating innate and metabolic pathways in homeostatic conditions are poorly understood. Here, we unveil a negative regulatory feedback loop involving the Stimulator of interferon genes (Sting) and the Fatty acid desaturase 2 (Fads2). At steady state, Sting regulates FA metabolism by repressing the activity of the Fads2 enzyme responsible for the desaturation of polyunsaturated FAs (PUFAs). Importantly, Sting activation increased Fads2 activity, while antagonizing Fads2 enhanced Sting activation, promoting the establishment of an anti-viral state. Remarkably, the cross-regulation between Sting and Fads2 is mediated by the cyclic GMP-AMP (cGAMP) Sting agonist and PUFAs. Indeed, we found that PUFAs inhibit Sting activation, while Sting agonists bind Fads2. Thus, our study identifies Sting as a master regulator of FA metabolism, and PUFAs as modulators of Sting-dependent inflammation. The interplay between Fads2 and Sting determines the fine-tuning of inflammatory responses, but comes at the expense of metabolic alterations, which are critical to consider in human pathologies associated with aberrant Sting activation.

## Introduction

The endoplasmic-reticulum (ER)-resident Stimulator of interferon genes (Sting) adaptor protein is central to the mounting of inflammatory responses in the presence of pathological nucleic acids, including dsDNA (Ishikawa et al., 2009). Indeed, aberrant dsDNA accumulation under stress conditions (Bai et al., 2017; King et al., 2017) or following pathogen infections (Gao et al., 2013a) can be detected by the cyclic GMP-AMP (cGAMP) synthase (cGAS) (Sun et al., 2013), that catalyzes the production of cGAMP (Ablasser et al., 2013; Gao et al., 2013b). This second messenger interacts with Sting (Zhang et al., 2013), promoting its activation through the recruitment of the Tank binding kinase 1 (Tbk1), resulting in phosphorylation-dependent activation of transcription factors, such as the Interferon regulatory factor 3 (Irf3) (Liu et al., 2015). This signaling pathway triggers the production of inflammatory cytokines and type I Interferons (IFNs) (Ishikawa et al., 2009). Dysregulations of Sting-associated signaling have been reported in a vast array of human pathologies. Yet, Sting function in absence of inflammatory challenge is unknown.

## Results

### Absence of STING Leads to Metabolic Improvements *In Vivo*

Because homeostasis requires tight regulation of metabolic and immune pathways (Brestoff and Artis, 2015; Buck et al., 2017), we questioned the impact of Sting ablation on metabolic parameters, at steady-state. Under normal diet, wild-type (WT) and Sting^-/-^ mice do not exhibit spontaneous inflammation (Figure 1A), nor differences in body weight (Figure 1B) and composition (Figure 1C). Yet, Sting^-/-^ mice present increased food intake (Figure 1D) and improved insulin-independent (Figure 1E and S1A) glucose management (Figure 1F), coupled to decreased hepatic gluconeogenesis (Figure 1G). Indirect calorimetry measurements showed that Sting^-/-^ mice consume more oxygen (Figure 1H) and present higher energy expenditure during the light phase (Figure 1I), in absence of change in circadian rhythm (Figure S1B) or spontaneous locomotor activity (Figure 1J). In addition, Sting^-/-^ mice, display increased thermogenesis (Figure 1K). Intriguingly, no significant change was measured in the expression of thermogenic program genes in the brown adipose tissue (Figure S1C), while *Uncoupling Protein 1* (*Ucp1*) and *Peroxisome proliferator-activated receptor-gamma coactivator-1alpha* (*Pgc-1α*) mRNA levels were increased in the visceral white adipose tissue (Figure 1L), reflecting the activation of browning pathways (Seale et al., 2007). Furthermore, the survival of Sting^-/-^ mice is increased as compared to WT mice under high fat diet (Figure 1M). Finally, metabolic phenotyping of cGAS-deficient mice and of conditional myeloid cell-specific Sting^--^ mice (Figure S1D-I) confirmed that metabolic alterations witnessed in Sting-deficient mice is independent of its canonical innate immune function. Therefore, absence of Sting is sufficient to cause global metabolic improvements *in vivo*.

**Figure 1:**
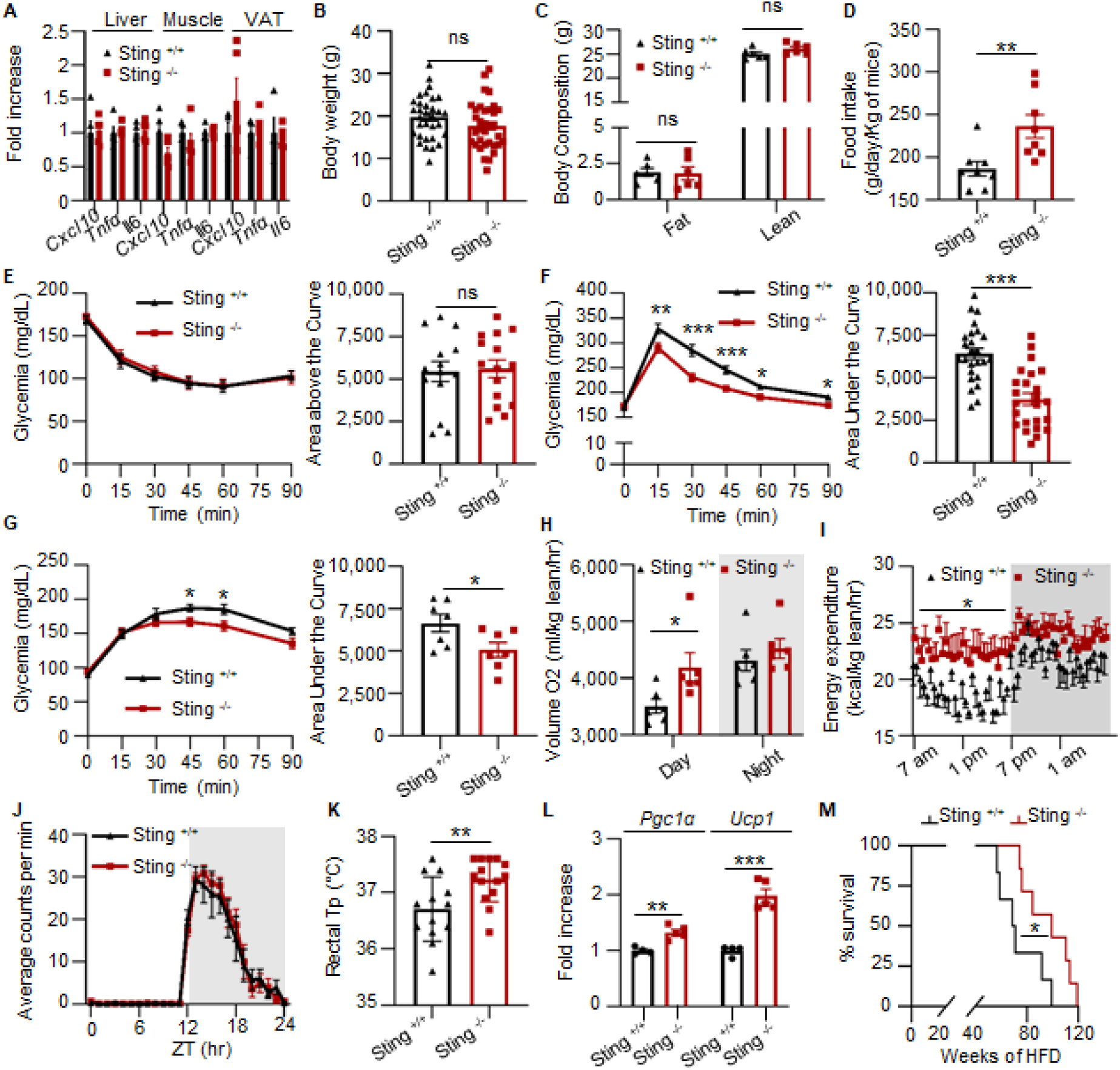
Sting Deficiency Leads to Global Metabolic Improvement. (A) *Cxcl10, Tnf-α* and *Il6* mRNA levels were measured in liver, muscle and visceral fat from Sting^+/+^ (n=5) and Sting^-/-^ (n=5) mice. (B) Body weight of Sting^+/+^ (n=32) and Sting^-/-^ (n=34) mice under normal diet. (C) Body composition of Sting^+/+^ (n=6) and Sting^-/-^ (n=6) mice was assessed by EchoMRI. (D) Food intake of Sting^+/+^ (n=8) and Sting^-/-^ (n=9) mice. (E) Insulin tolerance test (ITT) in Sting^+/+^ (n=14) and Sting^-/-^ (n=15) mice. Left panel: Glycemia (mg/dL) over time, following a bolus of Insulin. Right panel, area above the curve. (F) Glucose tolerance test (GTT) was performed in Sting^+/+^ (n=25) and Sting^-/-^ (n=24) mice. Left panel: Glycemia (mg/dL) over time, following a bolus of Glucose. Right panel, area under the curve (AUC). (G) Pyruvate tolerance test (PTT) was performed in Sting^+/+^ (n=7) and Sting^-/-^ (n=7) mice. Left panel: Glycemia (mg/dL) over time, following a bolus of Pyruvate. Right panel: AUC. (H) Oxygen consumption of Sting^+/+^ (n=6) and Sting^-/-^ (n=6) mice as determined in metabolic chambers. (I) Energy expenditure during day (white) and night (grey) was determined as in (H). P-value was determined by One-way Anova. (J) Daily profile of voluntary running-wheel activity of Sting^+/+^ (n=11) and Sting^-/-^ (n=10) mice. (K) Rectal temperature of Sting^+/+^ (n=13) and Sting^-/-^ (n=14) mice. (L) *Pgc1α* and *Ucp1* mRNA levels in the white adipose tissue from Sting^+/+^ (n=4) and Sting^-/-^ (n=5) mice. (M) Survival curve of Sting^+/+^ (n=6) and Sting^-/-^ (n=7) mice under high fat diet (HFD 60%) All graphs present means ± Standard Error of the Mean (SEM). P-value was determined by Student’s t-test, unless otherwise stated. ns: not significant, *: P < 0.05, **P: < 0.01, ***P: < 0.001.

### Sting interacts with and inhibits the Fatty Acid Desaturase 2

To identify the molecular mechanism through which Sting regulates metabolic homeostasis, we performed tandem-affinity purification of Flag- and HA-tagged Sting (F/HA-Sting) stably expressed in mouse embryonic fibroblasts (MEF) knockout of Sting (MEF^Sting-/-^). Immunopurified material was either silver-stained (Figure 2A) or analyzed by Mass spectrometry to identify Sting protein partners. Besides known Sting interactors, this approach revealed a large number of proteins involved in metabolic pathways (Table S1), notably including the RE-resident Fatty acid desaturase 2 (Fads2). Fads2 is the first, rate-limiting enzyme in the desaturation of linoleic acid [LA (18:2n-6, or Omega-6)] and α-linolenic acid [ALA (18:3n-3, Omega-3)] precursors (Nakamura and Nara, 2004), to generate polyunsaturated fatty acids (PUFAs) (Figure 2B). Precisely, Fads2 catalyses the desaturation of ALA into eicosapentaenoic acid (EPA), and subsequently into docosahexaenoic acid (DHA). Desaturation of LA into Dihomo-γ-linolenic acid (DGLA) also requires Fads2 (Nakamura and Nara, 2004), while further desaturation into arachidonic acid (AA) is catalysed by the Fatty acid desaturate 1 (Fads1) (Leonard et al., 2000). Dedicated enzymes further process these PUFAs into Oxylipins that influence numerous physiological processes (Gabbs et al., 2015). The interaction between Fads2 and Sting was verified by Western blot (WB) analysis of Flag-immunoprecipitated F/HA-Sting (Figure 2C). Conversely, Flag-immunoprecipitation of Flag-tagged Fads2 (F-Fads2) allowed co-immunoprecipitation of Sting, but not of Tbk1 (Figure 2D). Thus, we show that Fads2 is a protein partner of Sting, independently of Tbk1 and of pro-inflammatory stimulation.

**Figure 2:**
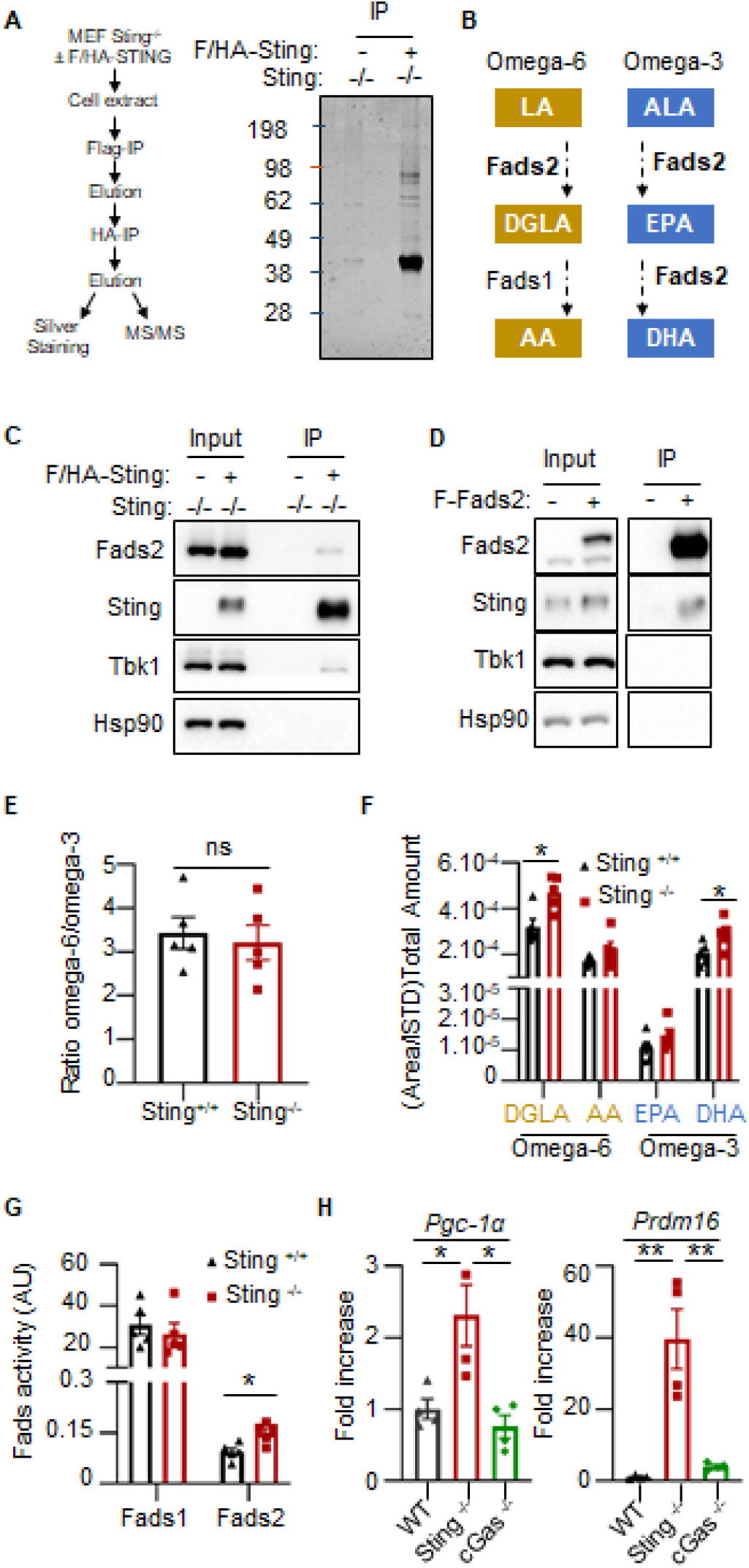
Sting Interacts with Fads2 and Modulates Polyunsaturated Fatty Acids Pools. (A) Left: Experimental scheme. Right: silver-staining of immunopurified Flag- and HA-tagged Sting (F/HA-Sting) separated on SDS-PAGE. Numbers on the left: molecular weight in kDa. (B) Schematic representation of the LA (Omega-6, Yellow) and ALA (Omega-3, Blue) fatty acids (FA) desaturation pathway, leading to the generation of polyunsaturated FA (PUFAs). ALA: α-linolenic acid, LA: linoleic acid, DGLA: Dihomo-γ -linolenic acid, AA: arachidonic acid, EPA: eicosapentaenoic acid, DHA: docosahexaenoic acid (C) Inputs and eluates from Flag-immunoprecipitated F/HA-Sting were analyzed by Western Blot (WB) using indicated antibodies. (D) Inputs and eluates from Flag-immunoprecipitated Flag-Fads2 were analyzed by WB using indicated antibodies. (E) Ratio between total Omega-6 and Omega-3 PUFAs and derivatives measured in Sting^+/+^ (n=5) and Sting^-/-^ (n=5) mice liver using LC-MS. (F) Sum of indicated PUFA and respective derivatives in samples analysed as in (E). (G) Fads1 and 2 enzyme activity was estimated by calculating the substrate/product ratio of PUFAs measured in (E). All graphs present means ± SEM. P-value was determined by Student’s t-test. ns: not significant, *: P < 0.05.

Because PUFAs are ligands of transcription factors that control thermogenesis (Fan et al., 2019), changes in PUFAs levels could be expected to promote the metabolic alterations observed in the absence of Sting (Figure 1). We used liquid chromatography coupled to mass spectrometry (LC-MS) to quantify PUFAs and their derivatives in liver and adipose tissue (AT) samples from WT and Sting^-/-^ mice. Partial least squares discriminant analysis (PLS-DA) of LC-MS data showed that WT and Sting^-/-^ liver and AT samples are significantly different (Figure S2A-B). Correlation analysis showed a shift in PUFA content, leading to accumulation of Omega-3 derivatives (Figure S2C-D). Although no significant shift in the total Omega-6/Omega-3 ratio was measured (Figure 2E), the total amount of derivatives from the main PUFAs families in mice liver samples showed that absence of Sting leads to significantly increased levels of DGLA and DHA derivatives (Figure 2F), coupled to increased Fads2 activity in Sting^-/-^ mice (Figure 2G). Similar increase in PUFAs deriving from Fads2 activity was measured in MEF^Sting-/-^ as compared to WT-MEFs, but not in MEF^cGAS-/-^ (Figure S2E-F). Importantly, mRNA levels of the *Pgc-1α* and *PR domain containing 16* (*Prdm16*) transcription factors involved in browning (Seale et al., 2007), were upregulation in Sting^-/-^, but not in cGAS^-/-^ MEFs (Figure 2H). This strongly supports that changes in PUFAs pools drive activation of thermogenesis program in absence of Sting.

### Sting Activation Promotes Fads2 activity

Because Sting activation leads to its degradation (Konno et al., 2013), we hypothesized that following Sting activation, the block of Fads2 activity would be alleviated. We used dsDNA transfection to activate Sting-dependent signalling (Figure 3A) and Sting degradation (Figure 3B), prior to analysis of PUFAs. We observed an increase in PUFAs deriving from Fads2 activity, including DHA (Figure 3C), accompanied by increased expression of *Pgc-1α* and *Prdm16* (Figure S3A), without significant shift in the Omega-6/Omega-3 balance (Figure 3D). Intriguingly, we also observed decreased Fads2 levels following Sting activation (Figure 3A), independently of type I IFN production (Figure S3B-C). This is in agreement with previous reports that Fads2 levels are decreased following its over-activation (Ralston et al., 2015). Of note, decreased Fads2 levels were also observed following infection with the Herpes Simplex Virus type I (HSV-1) DNA virus known to promote Sting activation (Figure S3D-F). Thus, acute activation-dependent decrease of Sting levels promotes Fads2 activity and metabolic reprogramming.

**Figure 3:**
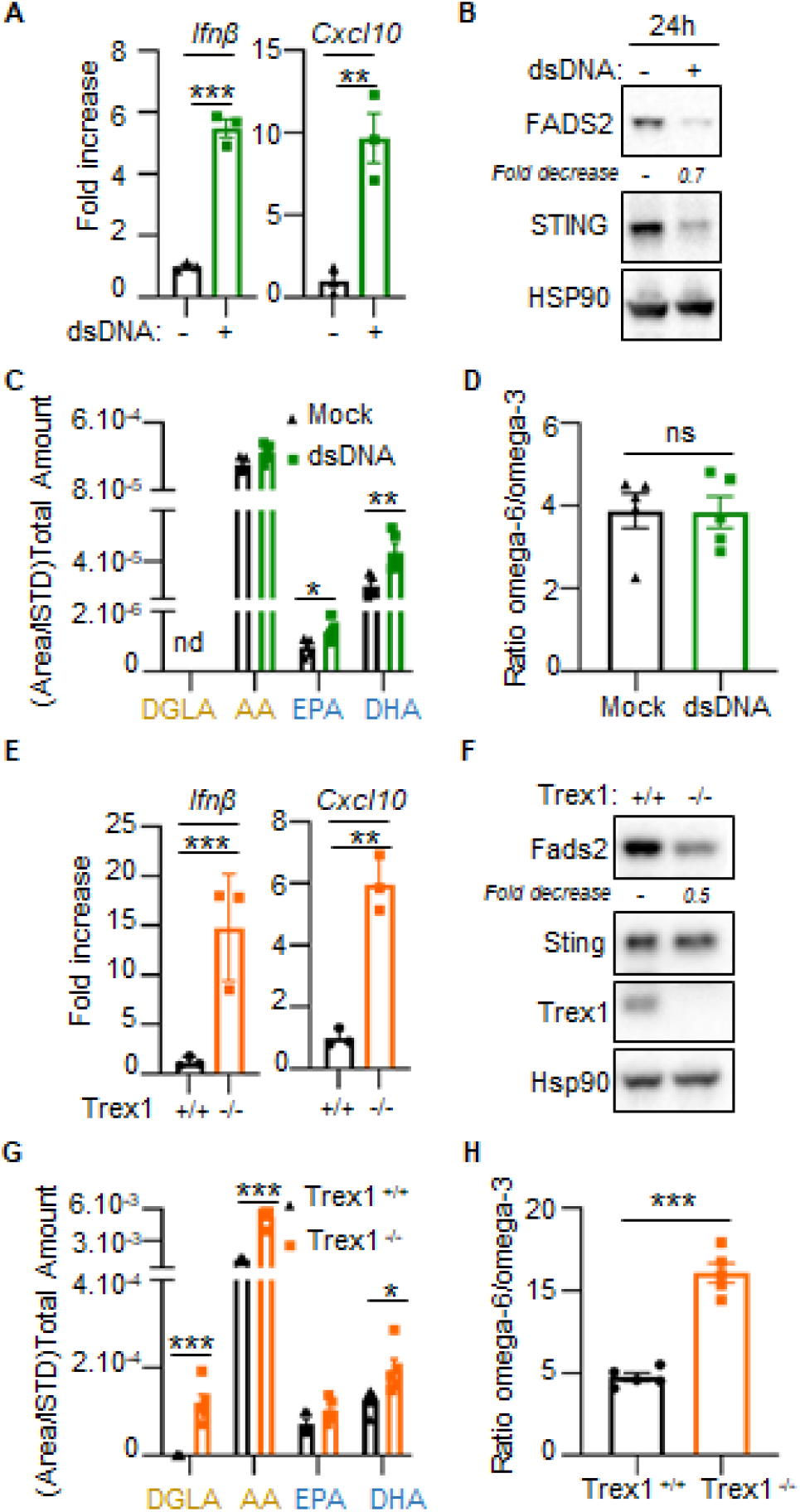
Activation-dependent Sting Degradation Promotes Fads2 Activity. (A) *Ifnβ* and *Cxcl10* mRNA levels in 293T cells transfected or not with dsDNA for 24h (n=3) (B) Whole cell extracts (WCE) of cells treated as in (A) were analyzed by WB using indicated antibodies. (C) Sum of indicated PUFA and derivatives in samples prepared as in (A). (D) Ratio between total Omega-6 and Omega-3 PUFAs and derivatives in samples from (C). (E) *Ifnβ* and *Cxcl10* mRNA levels in WT-MEF (n=3) and MEF^Trex1-/-^ (n=3). (F) WCE from WT-MEF or MEF^Trex1-/-^ were analyzed by WB using indicated antibodies. (G) Sum of indicated PUFAs and derivatives in WT-MEF or MEF^Trex1-/-^ (n=5). (H) Ratio between total Omega-6 and Omega-3 PUFAs and derivatives in samples from (G). All graphs present means ± SEM. P-value was determined by Student’s t-test. *: P < 0.05, **P: < 0.01, ***P: < 0.001.

To assess the impact of chronic Sting activation on Fads2 activity, we used MEF knockout of the Three prime exonuclease 1 (MEF^Trex1-/-^) that present with spontaneous chronic Sting activation (Ablasser et al., 2014) (Figure 3E). Absence of Trex1 correlates with decreased Fads2 protein levels, in absence of change in Sting protein levels (Figure 3F). Analysis of PUFAs and derivatives in WT and Trex1^-/-^ MEFs showed increased DGLA and DHA levels, indicating increased Fads2 activity (Figure 3G). However, we also measured increased levels of AA in absence of Trex1 (Figure 3G), together with an increased Omega-6/Omega-3 ratio (Figure 3H). While this supports our model whereby Sting activation promotes increased Fads2 activity, this also indicates that Sting degradation is not the sole parameter modulating Fads2 activity upon chronic Sting activation.

### cGAMP and PUFAs Orchestrate the Crosstalk Between Fads2 and Sting

Because Sting activation requires its interaction with dinucleotides, we hypothesized that the latter may participate to the control of Fads2 activation. We first tested this hypothesis *in silico* by docking cGAMP and the 5,6-dimethylxanthenone-4-acetic acid (DMXAA) Sting agonist into Fads2. To this aim, we used the resolved crystal of Sting in complex with cGAMP (PDB ID: 6WD4) and the molecular model of Fads2 as starting biological systems. This predicted that cGAMP and DMXAA can dock into Fads2, adopting similar conformation (Figure 4A, left and right panels), achieving analogous interactions and docking energies (Figure S4A and Table S2). We confirmed the interaction between cGAMP and Fads2 using *in vitro* binding assays (Figure 4B). In addition, we treated cGas^-/-^, Sting^-/-^, and WT-MEF with DMXAA, or performed dsDNA transfection to induce cGAMP production, prior to analysis of Fads2 protein levels. This showed that Fads2 levels are decreased in presence of DMXAA, regardless of the expression of cGas and Sting (Figure 4C), while dsDNA transfection-induced Fads2 degradation was reduced in the absence of cGas (Figure S4B). Thus, altogether these data show that Fads2 is a target of dinucleotides targeting Sting.

**Figure 4:**
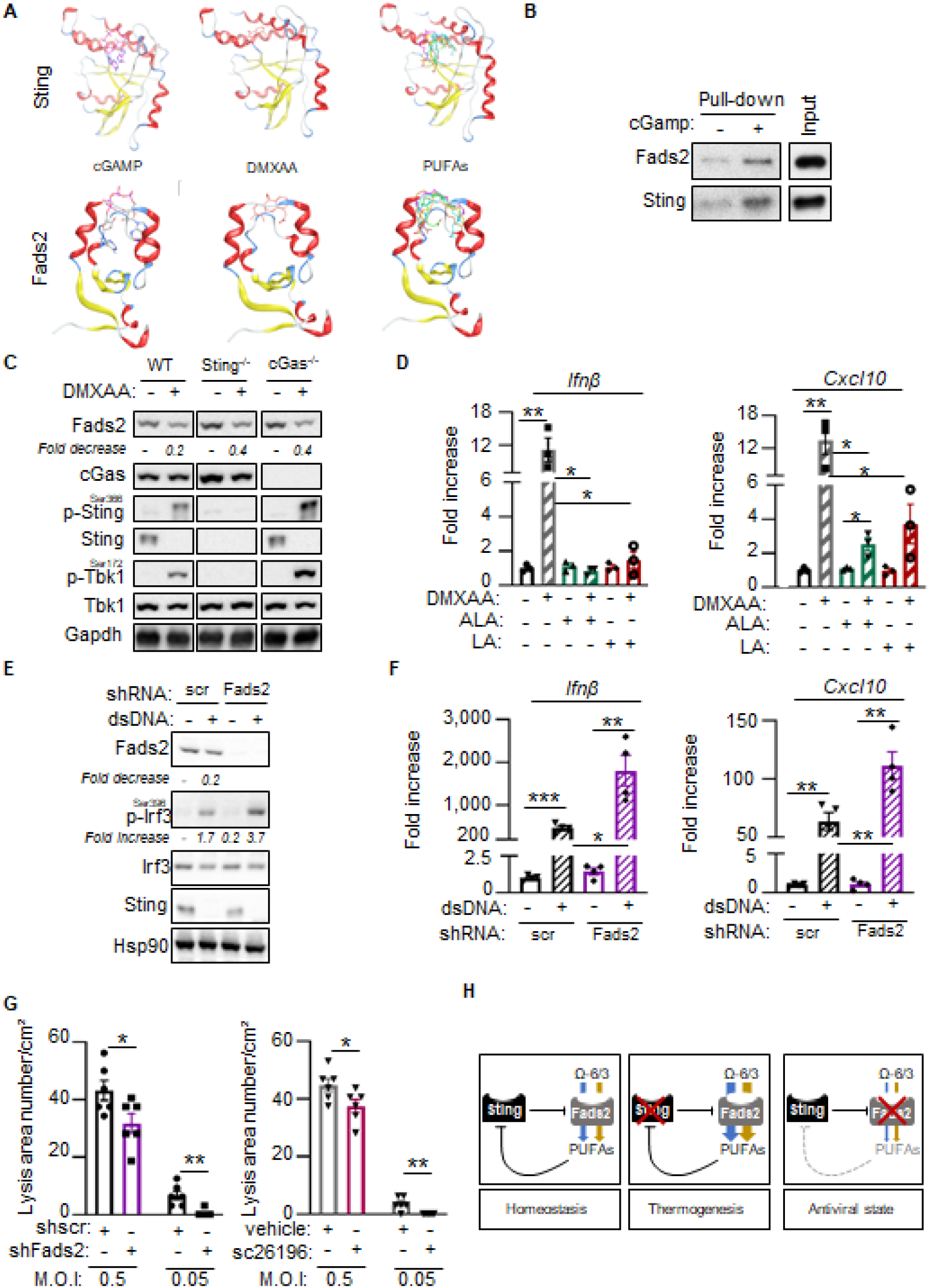
cGAMP and PUFAs Orchestrate the Crosstalk between Fads2 and Sting. (A) Molecular docking of cGAMP, DMXAA, and 6 PUFAs to STING (top) or FADS2 (bottom). Color coding for ligand: LA in blue, ALA in green, AA in orange, DHA in turquoise, DGLA in brown, and EPA in magenta. (B) Binding of Flag-purified F-Fads2 or F/HA-Sting to cGAMP was analysed by WB using anti-Sting or anti-Fads2 antibodies. (C) WCE from WT, Sting^-/-^, or cGas^-/-^ MEF stimulated or not with DMXAA for 2 h were analyzed by WB using indicated antibodies. (D) *Ifnβ* and *Cxcl10* mRNA levels in WT-MEF stimulated or not with DMXAA in combination or not with ALA or LA (n=3). (E) WCE from MEF expressing non-targeting (shScr) or Fads2-targetting (shFads2) shRNAs, transfected or not with dsDNA for 6 h, were analyzed by WB using indicated antibodies. (F) *Ifnβ* and *Cxcl10* mRNA levels in cells expressing Scramble (scr) or Fads2-targeting shRNAs after stimulation or not with dsDNA for 6 h (n=4). (G) WT-MEF were infected with HSV-KOS64 in presence or not of a Fads2 inhibitor (sc26196). Infection is presented as mean lysis area per cm^2^. (H) Schematic representation of the crosstalk between Sting and Fads2. All graphs are means ± SEM from at least 3 independent experiments. P-value was determined by Student’s t-test.*: P < 0.05, **: P < 0.01, ***: P < 0.001

Intriguingly, molecular docking analysis between PUFAs and Sting, predict that PUFAs can dock to the cGAMP-binding domain of Sting (Figure 4A, right panels). Indeed, cGAMP, DMXAA and PUFAs, adopt similar conformations in *in silico* docking experiments performed with Sting or Fads2 (Figure S4A), achieving analogous binding energies (Table S2). Furthermore, Sting and Fads2 showed strong structural similarity, both with regards to the 3D arrangement of the docking site anatomy, and in terms of hydrophobicity, electrostatics and solvent accessibility (Figure S4A). This suggested that PUFAs are potential ligands of Sting. We thus tested the impact of administrating ALA and LA precursors to cells prior to assessment of DMXAA-induced Sting activation. Both treatments decreased DMXAA-dependent expression of *Ifnβ* and of the *Cxcl10* interferon stimulated gene (ISG) (Figure 4D). Consistently, knock-down of Fads2, which caused a global decrease of Fads2 by-products (Figure S4C), led to increased *Ifnβ* and *Cxcl10* expression (Figure 4E-F), together with decreased expression of thermogenesis program genes (Figure S4D). Finally, antagonizing Fads2, using shRNAs or an inhibitor, led to decreased infection by HSV-1 (Figure 4G), consistent with the establishment of an antiviral state. Thus, we establish that PUFAs as inhibitors of Sting. Altogether, we show that Fads2 and Sting cooperate for the maintenance of global homeostasis, through fine-tuning of inflammatory and metabolic pathways (Figure 4H).

## Discussion

Altogether, we uncover a central role of Sting in the regulation of metabolic homeostasis, independently of its reported innate immune function. Indeed, we show that Sting inhibits Fads2 activity, thereby altering PUFAs pools. Absence of Sting thus increases the levels of Omega-3-derived PUFAs that in turn improve glucose handling (Sirtori and Galli, 2002), prevent obesity-associated glucose intolerance (Belchior et al., 2015; Derosa et al., 2016) and increase cardiovascular protection (Gonzalez-Periz et al., 2009). In agreement, we found that absence of Sting leads to better survival of mice under high fat diet. Furthermore, Omega-3-derived PUFAs can promote white adipose tissue browning (Fernández-Galilea et al., 2020), a process in which increased *Ucp1, Pgc-1α* and *Prdm16* expression is crucial (Ghandour et al., 2018). We found that absence of Sting promotes Fads2-dependent induction of these thermogenic factors, demonstrating that metabolic remodelling in absence of Sting requires Fads2. In addition, we show that Sting agonists, can directly interact with Fads2 and promote its degradation, establishing Fads2 as a direct target of cGAMP and DMXAA. This implies that in pathologies presenting with chronic Sting activation, Sting-dependent modulation of Fads2 activity may feed metabolic comorbidities such as dyslipidemia (Lira et al., 2014) or cachexia (Baazim et al., 2019). Furthermore, small molecules targeting Sting (Haag et al., 2018; Wu et al., 2020) can be expected to alter Fads2 activity and downstream production of inflammatory lipid mediators.

That PUFAs can inhibit Sting, reveals a previously unappreciated link between fatty acid metabolism and innate immune responses. Indeed, there are indications that diet intervention and modulation of Omega-3 or Omega-6 intake can impact immune and antiviral responses (DiNicolantonio and O’Keefe, 2018), although the involved molecular mechanism remaining poorly understood. We reveal that Fads2 activity and PUFAs can inhibit Sting activation. While his process may serve in the resolution of Sting-dependent inflammation, dietary habits that impact the substrates provided to Fads2 for desaturation (Galland, 2010; Tosi et al., 2014) may directly influence Sting activation and alter immune function (Sen et al., 2019). In addition, we show that modulating Fads2 activity impacts on the establishment of an antiviral state. This is in agreement with previous reports that PUFAs levels can both influence (Berra et al., 2017) and be influenced by HSV-1 infection (Zhang et al., 2020). Thus, we reveal a novel mechanism through which PUFAs contribute to the fine-tuning of inflammatory responses and identify Sting as a trigger of thermogenic program. Altogether, our findings offer unprecedented insight into the crosstalk between innate immune processes and metabolic regulation. Targeting this crosstalk in pathologies presenting with chronic inflammation, bears the promise to alleviate associated comorbidities.

## Acknowledgements

We thank M Benkirane, G Cavalli, J Déjardin and B de Massy for discussions and comments. We acknowledge: the SIRIC Montpellier Cancer Grant (INCa_Inserm_DGOS_12553), Metamus-RAM and iExplore-RAM animal facilities, the “Laboratoire de Mesures Physiques” of the University of Montpellier, for access to the Mass Spectrometry instruments, the MRI imaging facility, member of the national infrastructure France-BioImaging infrastructure supported by the French National Research Agency (ANR-10-INBS-04, “Investments for the future”) and Ross Tomaino from the “Taplin Mass Spectrometry Facility” of the Harvard Medical School, for Mass Spectrometry analysis. Work in NL’s laboratory is supported by the European Research Council (ERC-Stg CrIC: 637763, ERC-PoC DIM-CrIC: 893772), “LA LIGUE pour la recherche contre le cancer” and the “Agence Nationale de Recherche sur le SIDA et les Hépatites virales” (ANRS). HC is supported by a PhD fellowhip from” LA LIGUE pour la recherche contre le cancer”. CT was supported by Merck Sharp and Dohme Avenir (MSD-Avenir – GnoSTic) program, followed by an ANRS fellowship. JM is supported by a “Conventions Industrielles de Formation par la Recherche” (CIFRE) fellowship from the “Agence Nationale de Recherche Technologie” (ANRT). AS is supported by the ERC-PoC DIM-CrIC (893772). IKV was supported by the ERC-Stg CrIC (637763) followed by the Fondation pour la Recherche Médicale (ARF20170938586). Work in SRP’s laboratory is supported by the European Research Council (ERC-AdG ENVISION; 786602), The Novo Nordisk Foundation (NNF18OC0030274), and the Lundbeck Foundation (R198-2015-171; R268-2016-3927). Work in AT’s laboratory is supported by a SIRIC Montpellier Cancer Grant INCa_Inserm_DGOS_12553, the Foundation de France (grant no. 00078461) and a LabEx MabImprove Starting Grant. XB is supported by ANR GH-gen (ANR-18-CE14-0017).

## Author contributions

IKV and NL conceived the study and designed experiments. IKV, HC, AS, CT, LSR, ET, MS, JM, XB, DV, AT performed experiments. IKV, SRP, DV, AT and NL supervised the study. JL provided mice and mice tissues samples. IKV, DV, AT and NL analyzed data and prepared figures. IKV, DV, AT and NL wrote the manuscript. All authors read and approved the final version of the manuscript.

## Materials & Correspondence

All raw data and materials generated in the course of the present work are available upon request.

## STAR Methods

### Animals

Animal protocols were performed in accordance with French and European Animal Care Facility guidelines. All experiments were approved by the Animal Welfare and Ethical Review Body of Languedoc-Roussillon. Housing and experimental procedures were approved by the French Agriculture and Forestry Ministry (A34-172-13 & 15040-2018050214043878). Males from 8 to 14 weeks of age were used for this study. Mice Sting deficient (Tmem173<tm1Camb>) were provided by Pr. Lei Jin (Jin et al., 2013). cGAS deficient mice line was purchased from the EMMA consortium (Strasbourg, France). Tmem173<tm1Camb> mice (conditional ready) were crossed with mice expressing FLP recombinase to obtain Sting floxed mice. Specific Tmem173 knock out was obtained by crossing homozygous Tmem173fl/fl mice with transgenic LysM-Cre mice (gift from Michael Hahne, IGMM, Montpellier) expressing the Cre recombinase under the control of a myeloid gene promoter.

### Mouse studies

Body weight was measured at 8 weeks of age. Body composition (Fat and lean mass) was evaluated by quantitative nuclear magnetic resonance imaging (EchoMRI 3-in-1 system; Echo Medical Systems, Houston, Texas) before metabolic chambers experiments. Metabolic rates in mice were measured by indirect calorimetry using a Comprehensive Lab Animal Monitoring System (CLAMS, Columbus Instruments) as previously described (Vila et al., 2019). Mice were housed individually in metabolic chambers with free access to water and food for 2 days for acclimatization before animals were returned to metabolic chambers and monitored for the next day for oxygen consumption (VO2), carbon dioxide production (VCO2), and food intake. Energy expenditure was calculated using the formula energy expenditure = (3.815 + 1.232 VO2/VCO2) × VO2, and normalized to lean body mass. Body temperature was assessed in mice using a RET-3 rectal probe (Kent Scientific). ITTs and GTTs were performed as previously described (Vila et al., 2014). Briefly, mice were fasted for 6 h with free access to drinking water. For ITTs, insulin was administered intraperitoneally (0.4 mU/g of mice fed normal chow), and blood glucose was measured at various times after injection from the tip of the tail with a Glucometer (Accu-Chek, Roche). For GTTs, d-glucose was administered intraperitoneally (2 g/kg of mice), and blood glucose levels were monitored. At 15 min after glucose injection, blood was collected for insulin quantitation. Serum insulin concentrations were determined by ELISA (Mouse Ultrasensitive Insulin ELISA, ALPCO Diagnostics). For PTT, mice were fasted for 18 h, followed by intraperitoneal injection of pyruvate (2 g/kg of mice), and blood glucose levels were monitored. To measure spontaneous locomotion and circadian behaviour, mice were housed individually in cages equipped with a running wheel. Voluntary activity was measured as running wheel revolutions recorded in one-minute bins and analysed with the ClockLab software (Actimetrics). Circadian behaviour was accessed during a period of light/dark cycle and under constant darkness as previously described (Abitbol et al., 2017). HSV-1 brain infections in mice were conducted as previously described (Reinert et al., 2021).

### Diets

At the age of 8 weeks mice were fed with HFD (60% energy as fat, Safe Diet).

### RNA Extraction and Real-Time PCR

Total RNA from tissues was extracted with Trizol reagent (Invitrogen) and RNA extraction kit (Sigma). RNA was quantified with a Nanodrop spectrophotometer (ND-1000, Nanodrop Technologies). RNA (1-2 μg) was reverse transcribed using SuperScript IV reverse transcriptase (Invitrogen). Expression of specific mRNAs was determined with a LightCycler (Roche) using the SYBR green PCR master mix (Takara). Reactions were performed in duplicate, and relative amounts of cDNA were normalized to Glyceraldehyde-3-phosphate dehydrogenase (Gapdh) and/or heat shock protein 90 (Hsp90). RT-qPCR primer sequences are:

Human:

hGapdh: F-CTGGCGTCTTCACCACCATGG; R-CATCACGCCACAGTTTCCCGG;

hIfnβ: F-GAATGGGAGGCTTGAATACTGCCT;

R-TAGCAAAGATGTTCTGGAGCATCTC;

Mouse:

mGapdh: F-TTCACCACCATGGAGAAGGC; R-GGCATCGACTGTGGTCATGA;

mIfnβ: F-CTGCGTTCCTGCTGTGCTTCTCCA; R-TTCTCCGTCATCTCCATAGGGATC;

mCxcl10: F-ATGACGGGCCAGTGAGAATG; R-TCAACACGTGGGCAGGATAG;

mHsp90: F-GTCCGCCGTGTGTTCATCAT; R-GCACTTCTTGACGATGTTCTTGC;

mTnf-α: F-CTGTAGCCCACGTCGTAGC; R-TTGAGATCCATGCCGTTG;

mUcp-1: F-CCTGCCTCTCTCGGAAACAA;R-TGTAGGCTGCCCAATGAACA;

mIl-6: F-GACTTCCATCCAGTTGCCTTCT; R-TCCTCTCCGGACTTGTGAAGTA

mPgc1α: F-AAAGGATGCGCTCTCGTTCA; R-GGAATATGGTGATCGGGAACA

mPrdm16: F-CAGCACGGTGAAGCCATTC; R-GCGTCGATCCGCTTGTG

### Western Blot Analysis

Tissues were homogenized using a Fastprep apparatus (MP) in a buffer containing 50 mM Tris-HCl (pH 8.0), 150 mM NaCl, 1% NP40, 0.5% Sodium deoxycholate, 0.1% SDS, 10 μl/ml protease inhibitor, 10 μl/ml phosphatase I inhibitor, and 10 μl/ml phosphatase II inhibitor. Cells were lysed in 5 packed cell volume of TENTG-150 [20 mM tris-HCl (pH 7.4), 0.5 mM EDTA, 150 mM NaCl, 10 mM KCl, 0.5% Triton X-100, 1.5 mM MgCl2, and 10% glycerol, supplemented with 10 mM β-mercaptoethanol, 0.5 mM PMSF and 1x phosphatase inhibitor] for 30 min at 4°C. Tissue and cell lysates were centrifuged at 14,000 g for 30 min at 4°C, and supernatants were stored at −80°C. Solubilized proteins (20-30 μg) from tissues or cells were run on 10% or 12% SDS-PAGE gels (Invitrogen Novex Tris-glycine) transferred onto nitrocellulose membrane (Biorad Trans blot turbo) and incubated with primary antibodies. Primary antibodies used include: anti-phospho IRF3 (1:500; Cell Signaling 4D4G), anti-IRF3(1:1000; Cell Signaling D6I4C), anti-phospho TBK1 (1:1000; Cell Signaling D52C2), anti-TBK1 (1:1000; Cell Signaling D1B4), anti-STING (1:1000; Cell Signaling D2P2F), anti-phospho STING (1:1000; Cell Signaling D8F4W), anti-glyceraldehyde-phosphate dehydrogenase (GAPDH; 1:5000; Proteintech Europe 800004-1-Ig), anti-TREX-1 (1:250; Santa Cruz Biotechnology C-11 sc133112), mouse specific anti-cGAS (1:1000; Cell Signaling D3080), anti-HSP90 (1:1000; Cell Signaling C45G5), and anti-FADS2 (1:10 000; Invitrogen PA5-87765). All secondary antibodies (Cell Signaling) were used at 1:2000 dilution. Immunoreactive proteins were visualized by chemiluminescence (SuperSignal West Pico or femto Thermo Scientific).

### Cells and cell cultures

293T, T98G, WT-MEF, MEF-Sting^-/-^, MEF-cGas^-/-^ were maintained in DMEM supplemented with 10% Fetal Bovine Serum (FBS), 1% Penicillin/Streptomycin and 1% Glutamine. MEF-Sting^-/-^ overexpressing FLAG- and HA-tagged Sting (F/HA-Sting) were generated by transducing MEF-Sting^-/-^ with retroviral particles packaging the pOZ-F/HA-Sting construct and selection with puromycin. WT-MEF overexpressing FLAG-tagged Fads2 (Flag-Fads2) were generated by transducing WT-MEF with retroviral particles packaging the pOZ-Flag-Fads2 construct and selection with puromycin. The T98G IRF3 and control knockout cell line was generated using LenticrisprV2-GFP system (Plasmid #82416) and cell sorting using a BD FACS Melody. Pooled cells were then amplified and IRF3 invalidation verified by Western Blot analysis.

### Cell treatment and transfection

Cells were transfected with JetPrime transfection reagent (Polyplus) at 1:2 ratio with various nucleic acids at 2 μg/ml. DMXAA (Invivogen) was used at 100 or 200 μM in Opti-MEM (Gibco). Sc26196 (Santa Cruz) was used at 2 μM.

### HSV-KOS64 amplification

HSV KOS-64-GFP strain was a gift from S. Paludan. The virus was amplified in Vero cells. Briefly, Vero cells were plated in T175 and infected with HSV KOS-64-GFP virus during 30 min. Media was subsequently replaced and cells were collected 72 h after infection for viral extraction using 3 freeze-thaw cycles. Two centrifugations steps were performed and concentrated virus was resuspended before storage at −80°C. 10^3^ cells were seeded in 96-wells plate for lysis area number or 2.10^5^ 6-wells plate for western blot 16h prior infection. Cells were infected with HSV KOS-64-GFP for 90min on presence or not of Fads2 inhibitor (2μM). Medium was replaced with DMEM/Human serum 2% medium for 16h. For lysis area number assessment, medium was replaced with DMEM/FBS 10% medium for 32h.

### Immunoprecipitation and mass spectrometry analysis

MEF-Sting^-/-^ overexpressing F/HA-Sting were lysed in 5 packed cell volume of TENTG-150. The first immunoprecipitation used an anti-FLAG antibody, followed by the elution using an excess of FLAG peptide. Eluates were subsequently used as input material for immunoprecipitation using an anti-HA antibody. Sting protein partners were eluted using an excess of HA peptide. Part of the FLAG and HA immunoprecipitated material was silver-stained and the remainder Coomassie-stained. Portions of the Coomassie-stained gel were excised and analyzed by Mass Spectrometry.

### In vitro pull-down using biotinylated cGAMP

Pull-down was carried out using 30 μl (0.3mg) of MyOne Streptavidin C1 Dynabeads per condition. An excess of Biotin or cGAMP was coupled to beads according to the manufacturer’s instructions. 30 μl of Flag-immunoprecipitated Fads2 or Sting was incubated with beads, on ice for 30 min in low-binding tubes (Axygen). Three consecutive washes were performed in 20 mM tris-HCl (pH 7.4), 10mM KCl, 0.5% Triton, 150 mM NaCl, 10% glycerol, 1.5 mM MgCl2 and 10 mM β-mercaptoethanol, and 0.5 mM PMSF. Tubes were changed at first and last washes. Bound material was eluted in 30 μl of Laemmli buffer.

### RNA interference

shRNA targeting Fads2 (Clone ID: NM_019699.1-487s1c1) and scramble (SHC016) were obtained from Sigma-Aldrich. shRNA-expressing lentiviral particles were produced by co-transfection of 2 × 10^6^ 293T cells with 5 μg of shFads2, 5 μg of psPAX2 (Gag-Pol), and 1 μg of pMD2G (Env), using the standard calcium-phosphate transfection protocol. Viral particles were harvested 48 h after transfection, filtered with 0.45 μM filters, and used for transduction. For knockdown of Fads2, 10^6^ MEF cells were seeded 24 h before transduction. Medium was replaced 10 h after transduction, and 1.5μg/ml puromycin selection was performed 72 h later.

### Measurement of PUFAs and oxilipins in biological samples

Snap-frozen tissues samples or 3× 10^6^ cells were crushed and solubilized (sonicated) in 1 mL of methanol (Wako, Tokyo, Japan). The samples were then spiked with following 16 internal standards (at 30 μM final concentration): tetranor-PGEM-d6, 6-keto-PGF1a-d4, TXB2-d4, PGF2a-d4, PGE2-d4, PGD2-d4, LTC4-d5, LTB4-d4, 15-HETE-d8, 12-HETE-d8, 5-HETE-d8, PAF-d4, OEA-d4, EPA-d5, DHA-d5, DHA-d5 and AA-d8 (all from Cayman Chemicals, Ann Arbor, MI, USA). The samples were then sonicated for 10 s, then vortexed for 2 min and further incubated for 2h at 4°C. Next, the samples were centrifugated at 15.000g for 10 min, the supernatant was removed and diluted with 4 mL of 0.1 % formic acid. The mix was briefly vortexed and loaded on the solid phase extraction (SPE) column Strata-X (Phenomenex, Torrance, CA, USA) in 1 ml steps. Before loading the sample, the SPE column was washed with 1 mL methanol and equilibrated with 2 mL of 0.1% formic acid. Following sample loading, the SPE column was washed with 1 mL 0.1% formic acid, 1 mL 15% ethanol and the compounds were eluted in 250 μL methanol. The eluate was lyophilized and then solubilized in 20 μL methanol, of which 5 μL sample were injected on UPLC coupled to triple-quadrupole MS (LCMS-8050). The sample was separated using analytical column Kinetex C8 (Cat. # 00F-4497-AN; Phenomenex), mobile phase A 0.1% formic acid (Sigma Aldrich) in water and mobile phase B acetonitrile (Sigma Aldrich). The flow rate was set at 0.4 mL/min, column oven temperature at 40°C. The gradient was: 0 min 90% A, 5 min 75% A, 10 min 65% A, 20 min 25% A, 20.1 – 25 min 5% A, 25.1 min 90% A. MS setting, data acquisition and data analysis were performed according to manufacturer instructions for analyzing Lipid Mediators version 2.0 (Cat. # 225-24873A, Shimadzu).

### LC-MS data analysis

Following visual inspection, integration and calculation of peak surface area for all identified compounds, the data were normalized using the internal standards. As outlined in the Lipid Mediators version 2.0 manual, individual compounds were repartitioned in 17 different groups, which were then normalized with 17 internal standards outlined above. Next, for each sample we calculated total metabolite load (TML), which was equal to the sum of individual metabolites measured in the respective sample. For TML calculation, missing values for individual metabolites were imputed by assigning the least possible peak area found in the dataset. The imputation was based on the hypothesis that as long as a given metabolite was present in some of the samples, its absence in other samples was due to the limit of detection of the analysis. All values for all metabolites within a sample were normalized with respect to TML. Once the normalized quantities were calculated, the data were imported in R (4.0.2) [R Core Team (2020)] and analyzed using the MetaboAnalyst package (4.0) (Chong et al., 2019).

### Homology Modelling

The homology modelling of the FADS2 was performed using the Molecular Operating Environment (MOE) Suite (Warde-Farley et al., 2010). The 3OZZ RCSB entry was used as template, which is the crystal structure of the *Bos taurus* cytochrome b5 core-swap mutant. The 3D models were subsequently energetically optimized the AMBER10 forcefield as it is implemented in the MOE Suite. Finally, all 3D models were assessed for their folding via the protein and geometry check of MOE Suite.

### Molecular Docking

The docking module of MOE was used for the docking of the 6 PUFAs or cGAMP/DMXAA to STING and the FADS2 model. The 6 PUFAs that were used were namely: linoleic acid (LA), α-linolanic acid (ALA), dihomo-γ-linolenic acid (DGLA), arachidonic acid (AA), docosahexaenoic acid (DHA) and eicosapentaenoic acid (EPA). A fast Fourier transformation (FFT) pipeline is utilized by MOE for the docking experiment. The overall score is influenced by the model’s packing, electrostatic, solvation and hydrophobic energies. Transient complexes of proteins are kept in a local database and their contact propensities are statistically used for docking. The top hits of the docking experiment were energetically optimized using energy minimization pipelines to relieve the models from any residual geometrical strain. Finally, the Drugster suite was used to perform a final and rapid energy minimization step using AMBER99 forcefield (Vilar et al., 2008), while solvated using an implicit Generalized Born (GB) water model.

### Molecular Dynamics

The interaction pattern and overall fold of the final complexes of each one of the 6 PUFAs or cGAMP/DMXAA to either STING and FADS2 model, were subjected to exhaustive molecular dynamics simulations using the DrugOn suite ^9^. Molecular dynamics simulations were executed in an explicitly SPC water solvated periodic cube system. Counterions were used as required to neutralize the molecular system. Each biological system was subjected to a hundred nanoseconds (100 ns) of molecular dynamics at 300K and at 1 fs step size. The molecular trajectory of each simulation was then imported into a local database for further analysis (Vlachakis et al., 2013).

## Supplementary Information

### Supplementary figure legends

**Figure S1: Sting Deficiency Leads to Global Metabolic Improvement**

(A) Insulinemia was measured 15 min after the injection of glucose during a glucose tolerance test (GTT) in Sting^+/+^ (n=6) and Sting^-/-^ (n=7) mice,

(B) Representative actograms of wheel activity measured under a 12 h light / 12 h dark schedule (yellow), and under constant darkness (blue) in Sting^+/+^ (n=11) and Sting^-/-^ (n=10) mice,

(C) *Ucp1* mRNA levels were measured in the brown adipose tissue from Sting^+/+^ (n=5) and Sting^-/-^ (n=6) mice,

(D) Body weight of cGas^+/+^ (n=6) and cGas^-/-^ (n=7) mice fed normal chow diet at 8 weeks age.

(E) GTT was performed in cGas^+/+^ (n=5) and cGas^-/-^ (n=5) mice,

(F) Rectal temperature was measured in cGas^+/+^ (n=6) and cGas^-/-^ (n=7) mice,

(G) Body weight of Sting-Flox^+/+^ (n=7) and Sting-Flox^LysM^ (n=8) mice fed normal chow diet at 8 weeks of age,

(H) GTT was performed in Sting-Flox^+/+^ (n=7) and Sting-Flox^LysM^ (n = 6) mice,

(I) Rectal temperature of Sting-Flox^+/+^ (n=9) and Sting-Flox^LysM^ (n=8) mice.

All graph present means ± SEM. P-values were determined by Student’s t-test. ns: not significant.

**Figure S2: Sting Interacts with Fads2 and Modulates Polyunsaturated Fatty Acids Pools**

(A) Partial least squares discriminant analysis (PLS-DA) of liver samples from Sting^+/+^ (pink) and Sting^-/-^ (green) mice in which PUFAs and derivatives were measured by LC-MS (n=5 mice per group),

(B) Partial least squares discriminant analysis (PLS-DA) of white visceral adipose tissue samples from Sting^+/+^ (pink) and Sting^-/-^ (green) mice in which PUFAs and derivatives were measured by LC-MS (n=3 mice per group),

(C) Correlation plots of PUFAs and derivatives measured in samples from (A),

(D) Correlation plots of PUFAs and derivatives measured in samples from (B),

(E) Ratio between total Omega-6 and Omega-3 PUFAs and derivatives in WT, Sting-/-, or cGas-/- MEF samples,

(F) PUFAs and derivatives were measured in WT, Sting-/-, or cGas-/- MEF using LC-MS. Graph presents the sum of indicated PUFA and respective derivatives.

In panels (C) and (D) the side bars indicate in green: derivatives of ALA (Omega-3), in red: derivatives from LA (Omega-6), in black: PUFAs that do not vary across the samples, in yellow: PUFAs and derivatives increased by at least 30%, and in blue: PUFAs decreased by at least 30%.

All graph present means ± SEM. P-values were determined by Student’s t-test. ns: not significant. *: P < 0.05, **P: < 0.01, ***P: < 0.001.

**Figure S3: Activation-dependent Sting Degradation Promotes Fads2 Activity**

(A) *Pgc1α* and *Prdm16* mRNA levels in WT-MEFs cells after stimulation or not with dsDNA (2μg) for 6 h (n=4),

(B) Whole cell extracts (WCE) from control (CTL) or IRF3 knockout T98G cell lines stimulated or not for 6 h with dsDNA were analyzed by WB using indicated antibodies,

(C) *Ifnβ* mRNA levels from samples treated as in (B) were analyzed by RT-qPCR. Graph presents mean fold induction ± SEM, as compared to unstimulated control cells (n=3),

(D) WCE from WT-MEFs, infected or not for 16 h with HSV-KOS64, were analyzed by WB using indicated antibodies. Multiplicity of infection (MOI) scale was from 1.5×10-4 to 1.5,

(E) Left: Representative mice brain samples infected or not for 5 days with the McKrae HSV-1 strain analyzed by WB using indicated antibodies. Right: Quantification of Fads2 protein levels in mice brain extracts following 5 days of infection or not with McKrae HSV1 (n=4),

(F) *Ifnβ* mRNA levels were quantified from mice brain samples treated as in (C)

All graph present means ± SEM. P-values were determined by Student’s t-test. ns: not significant. *: P < 0.05, **P: < 0.01, ***P: < 0.001.

**Figure S4: cGAMP and PUFAs Orchestrate the Crosstalk between Fads2 and Sting**

(A) PUFAs interact with the cGAMP-binding domain of Sting. 2D molecular interaction maps for cGAMP, DMXAA and the 6 PUFAs docked into STING (left column) and FADS2 (right column). LA: linoleic acid, DGLA: Dihomo-γ -linolenic acid, AA: arachidonic acid, ALA: α-linolenic acid, EPA: eicosapentaenoic acid, DHA: docosahexaenoic acid,

(B) WCE from WT, Sting-/-, or cGas-/- MEF stimulated or not dsDNA (2μg) for 6h (left) or 24 h (right) were analyzed by WB using indicated antibodies,

(C) PUFAs measured in cells expressing Scramble (scr) or Fads2-targeting shRNAs using LC-MS. Graph presents indicated PUFA,

(D) *Prdm16* mRNA levels in cells expressing Scramble (scr) or Fads2-targeting shRNAs after stimulation or not with dsDNA for 6 h (n=4).

All graph present means ± SEM. P-values were determined by Student’s t-test. ns: not significant. *: P < 0.05, **P: < 0.01.

### Supplementary Tables

**Table S1.** Major protein partners of Sting identified by Mass Spectrometry. * Unique peptides identified in Mock IP. **: Unique peptides identified in F/HA-Sting IP. MW is in kDa. IP: immunoprecipitation.

| Uniprot Reference | Gene Symbol | MW | Mock | IP |
| --- | --- | --- | --- | --- |
| sp\|Q9WUN2\|TBK1_MOUSE | Tbk1 | 83.37 | 0 | 8 |
| sp\|Q9Z0R9\|FADS2_MOUSE | Fads2 | 52.35 | 0 | 10 |
| sp\|Q920L1\|FADS1_MOUSE | Fads1 | 52.29 | 0 | 6 |
| sp\|Q9D6K9\|CERS5_MOUSE | Cers5 | 48.13 | 0 | 5 |
| sp\|Q9CY27\|TECR_MOUSE | Tecr | 36.07 | 0 | 5 |
| sp\|Q8VCH6\|DHC24_MOUSE | Dhcr24 | 60.07 | 0 | 4 |
| sp\|Q6NVG1\|LPCT4_MOUSE | Lpcat4 | 57.11 | 0 | 4 |
| sp\|Q8K2C9\|HACD3_MOUSE | Hacd3 | 43.10 | 0 | 2 |
| sp\|P47740\|AL3A2_MOUSE | Aldh3a2 | 53.94 | 0 | 2 |

**Table S2.** Interaction energies of the docked cGAMP, DMXAA and the 6 PUFAs on Sting and Fads2 proteins. Energies are in Kcal/mol.

|  | STING | FADS2 |
| --- | --- | --- |
| cGAMP | -34.925 | -7.269 |
| DMXAA | -24.400 | -51.661 |
| linoleic acid (LA) | -39.483 | -57.098 |
| dihomo-γ-linolenic acid (DGLA) | -13.064 | -47.331 |
| arachidonic acid (AA) | -42.827 | -58.912 |
| α-linolenic acid (ALA) | -43.658 | -38.473 |
| eicosapentaenoic acid (EPA) | -24.926 | -41.078 |
| docosahexaenoic acid (DHA) | -53.608 | -57.339 |

## References

Abitbol, K., Debiesse, S., Molino, F., Mesirca, P., Bidaud, I., Minami, Y., Mangoni, M.E., Yagita, K., Mollard, P., and Bonnefont, X. (2017). Clock-dependent and system-driven oscillators interact in the suprachiasmatic nuclei to pace mammalian circadian rhythms. PLoS One 12, e0187001.

Ablasser, A., Goldeck, M., Cavlar, T., Deimling, T., Witte, G., Röhl, I., Hopfner, K.P., Ludwig, J., and Hornung, V. (2013). cGAS produces a 2’-5’-linked cyclic dinucleotide second messenger that activates STING. Nature 498, 380–384.

Ablasser, A., Hemmerling, I., Schmid-Burgk, J.L., Behrendt, R., Roers, A., and Hornung, V. (2014). TREX1 deficiency triggers cell-autonomous immunity in a cGAS-dependent manner. Journal of immunology (Baltimore, Md.: 1950) 192, 5993–5997.

Baazim, H., Schweiger, M., Moschinger, M., Xu, H., Scherer, T., Popa, A., Gallage, S., Ali, A., Khamina, K., Kosack, L., et al. (2019). CD8(+) T cells induce cachexia during chronic viral infection. Nature immunology 20, 701–710.

Bai, J., Cervantes, C., Liu, J., He, S., Zhou, H., Zhang, B., Cai, H., Yin, D., Hu, D., Li, Z., et al. (2017). DsbA-L prevents obesity-induced inflammation and insulin resistance by suppressing the mtDNA release-activated cGAS-cGAMP-STING pathway. Proceedings of the National Academy of Sciences of the United States of America 114, 12196–12201.

Belchior, T., Paschoal, V.A., Magdalon, J., Chimin, P., Farias, T.M., Chaves-Filho, A.B., Gorjao, R., St-Pierre, P., Miyamoto, S., Kang, J.X., et al. (2015). Omega-3 fatty acids protect from diet-induced obesity, glucose intolerance, and adipose tissue inflammation through PPARgamma-dependent and PPARgamma-independent actions. Mol Nutr Food Res 59, 957–967.

Berra, A., Tau, J., Zapata, G., and Chiaradia, P. (2017). Effects of PUFAs in a Mouse Model of HSV-1 Chorioretinitis. Ocul Immunol Inflamm 25, 844–854.

Brestoff, J.R., and Artis, D. (2015). Immune regulation of metabolic homeostasis in health and disease. Cell 161, 146–160.

Buck, M.D., Sowell, R.T., Kaech, S.M., and Pearce, E.L. (2017). Metabolic Instruction of Immunity. Cell 169, 570–586.

Chong, J., Wishart, D.S., and Xia, J. (2019). Using MetaboAnalyst 4.0 for Comprehensive and Integrative Metabolomics Data Analysis. Curr Protoc Bioinformatics 68, e86.

Derosa, G., Cicero, A.F., D’Angelo, A., Borghi, C., and Maffioli, P. (2016). Effects of n-3 pufas on fasting plasma glucose and insulin resistance in patients with impaired fasting glucose or impaired glucose tolerance. Biofactors 42, 316–322.

DiNicolantonio, J.J., and O’Keefe, J.H. (2018). Importance of maintaining a low omega-6/omega-3 ratio for reducing inflammation. Open Heart 5, e000946.

Fan, R., Koehler, K., and Chung, S. (2019). Adaptive thermogenesis by dietary n-3 polyunsaturated fatty acids: Emerging evidence and mechanisms. Biochim Biophys Acta Mol Cell Biol Lipids 1864, 59–70.

Fernández-Galilea, M., Félix-Soriano, E., Colón-Mesa, I., Escoté, X., and Moreno-Aliaga, M.J. (2020). Omega-3 fatty acids as regulators of brown/beige adipose tissue: from mechanisms to therapeutic potential. Journal of physiology and biochemistry 76, 251–267.

Gabbs, M., Leng, S., Devassy, J.G., Monirujjaman, M., and Aukema, H.M. (2015). Advances in Our Understanding of Oxylipins Derived from Dietary PUFAs. Advances in nutrition (Bethesda, Md.) 6, 513–540.

Galland, L. (2010). Diet and inflammation. Nutrition in clinical practice: official publication of the American Society for Parenteral and Enteral Nutrition 25, 634–640.

Gao, D., Wu, J., Wu, Y.T., Du, F., Aroh, C., Yan, N., Sun, L., and Chen, Z.J. (2013a). Cyclic GMP-AMP synthase is an innate immune sensor of HIV and other retroviruses. Science (New York, N.Y.) 341, 903–906.

Gao, P., Ascano, M., Wu, Y., Barchet, W., Gaffney, B.L., Zillinger, T., Serganov, A.A., Liu, Y., Jones, R.A., Hartmann, G., et al. (2013b). Cyclic [G(2’,5’)pA(3’,5’)p] is the metazoan second messenger produced by DNA-activated cyclic GMP-AMP synthase. Cell 153, 1094–1107.

Ghandour, R.A., Colson, C., Giroud, M., Maurer, S., Rekima, S., Ailhaud, G., Klingenspor, M., Amri, E.Z., and Pisani, D.F. (2018). Impact of dietary omega3 polyunsaturated fatty acid supplementation on brown and brite adipocyte function. J Lipid Res 59, 452–461.

Gonzalez-Periz, A., Horrillo, R., Ferre, N., Gronert, K., Dong, B., Moran-Salvador, E., Titos, E., Martinez-Clemente, M., Lopez-Parra, M., Arroyo, V., et al. (2009). Obesity-induced insulin resistance and hepatic steatosis are alleviated by omega-3 fatty acids: a role for resolvins and protectins. FASEB J 23, 1946–1957.

Haag, S.M., Gulen, M.F., Reymond, L., Gibelin, A., Abrami, L., Decout, A., Heymann, M., van der Goot, F.G., Turcatti, G., Behrendt, R., et al. (2018). Targeting STING with covalent small-molecule inhibitors. Nature 559, 269–273.

Ishikawa, H., Ma, Z., and Barber, G.N. (2009). STING regulates intracellular DNA-mediated, type I interferon-dependent innate immunity. Nature 461, 788–792.

Jin, L., Getahun, A., Knowles, H.M., Mogan, J., Akerlund, L.J., Packard, T.A., Perraud, A.L., and Cambier, J.C. (2013). STING/MPYS mediates host defense against Listeria monocytogenes infection by regulating Ly6C(hi) monocyte migration. Journal of immunology (Baltimore, Md.: 1950) 190, 2835–2843.

King, K.R., Aguirre, A.D., Ye, Y.X., Sun, Y., Roh, J.D., Ng, R.P., Jr., Kohler, R.H., Arlauckas, S.P., Iwamoto, Y., Savol, A., et al. (2017). IRF3 and type I interferons fuel a fatal response to myocardial infarction. Nature medicine 23, 1481–1487.

Konno, H., Konno, K., and Barber, G.N. (2013). Cyclic dinucleotides trigger ULK1 (ATG1) phosphorylation of STING to prevent sustained innate immune signaling. Cell 155, 688–698.

Leonard, A.E., Kelder, B., Bobik, E.G., Chuang, L.T., Parker-Barnes, J.M., Thurmond, J.M., Kroeger, P.E., Kopchick, J.J., Huang, Y.S., and Mukerji, P. (2000). cDNA cloning and characterization of human Delta5-desaturase involved in the biosynthesis of arachidonic acid. The Biochemical journal 347 Pt 3, 719–724.

Lira, F.S., Rosa Neto, J.C., Antunes, B.M., and Fernandes, R.A. (2014). The relationship between inflammation, dyslipidemia and physical exercise: from the epidemiological to molecular approach. Curr Diabetes Rev 10, 391–396.

Liu, S., Cai, X., Wu, J., Cong, Q., Chen, X., Li, T., Du, F., Ren, J., Wu, Y.T., Grishin, N.V., et al. (2015). Phosphorylation of innate immune adaptor proteins MAVS, STING, and TRIF induces IRF3 activation. Science (New York, N.Y.) 347, aaa2630.

Nakamura, M.T., and Nara, T.Y. (2004). Structure, function, and dietary regulation of delta6, delta5, and delta9 desaturases. Annual review of nutrition 24, 345–376.

Ralston, J.C., Matravadia, S., Gaudio, N., Holloway, G.P., and Mutch, D.M. (2015). Polyunsaturated fatty acid regulation of adipocyte FADS1 and FADS2 expression and function. Obesity (Silver Spring, Md.) 23, 725–728.

Reinert, L.S., Rashidi, A.S., Tran, D.N., Katzilieris-Petras, G., Hvidt, A.K., Gohr, M., Fruhwurth, S., Bodda, C., Thomsen, M.K., Vendelbo, M.H., et al. (2021). Brain immune cells undergo cGAS/STING-dependent apoptosis during herpes simplex virus type 1 infection to limit type I IFN production. J Clin Invest 131.

Seale, P., Kajimura, S., Yang, W., Chin, S., Rohas, L.M., Uldry, M., Tavernier, G., Langin, D., and Spiegelman, B.M. (2007). Transcriptional control of brown fat determination by PRDM16. Cell Metab 6, 38–54.

Sen, T., Rodriguez, B.L., Chen, L., Corte, C.M.D., Morikawa, N., Fujimoto, J., Cristea, S., Nguyen, T., Diao, L., Li, L., et al. (2019). Targeting DNA Damage Response Promotes Antitumor Immunity through STING-Mediated T-cell Activation in Small Cell Lung Cancer. Cancer Discov 9, 646–661.

Sirtori, C.R., and Galli, C. (2002). N-3 fatty acids and diabetes. Biomed Pharmacother 56, 397–406.

Sun, L., Wu, J., Du, F., Chen, X., and Chen, Z.J. (2013). Cyclic GMP-AMP synthase is a cytosolic DNA sensor that activates the type I interferon pathway. Science (New York, N.Y.) 339, 786–791.

Tosi, F., Sartori, F., Guarini, P., Olivieri, O., and Martinelli, N. (2014). Delta-5 and delta-6 desaturases: crucial enzymes in polyunsaturated fatty acid-related pathways with pleiotropic influences in health and disease. Advances in experimental medicine and biology 824, 61–81.

Vila, I.K., Badin, P.M., Marques, M.A., Monbrun, L., Lefort, C., Mir, L., Louche, K., Bourlier, V., Roussel, B., Gui, P., et al. (2014). Immune cell Toll-like receptor 4 mediates the development of obesity- and endotoxemia-associated adipose tissue fibrosis. Cell Rep 7, 1116–1129.

Vila, I.K., Park, M.K., Setijono, S.R., Yao, Y., Kim, H., Badin, P.M., Choi, S., Narkar, V., Choi, S.W., Chung, J., et al. (2019). A muscle-specific UBE2O/AMPKalpha2 axis promotes insulin resistance and metabolic syndrome in obesity. JCI Insight 4.

Vilar, S., Cozza, G., and Moro, S. (2008). Medicinal chemistry and the molecular operating environment (MOE): application of QSAR and molecular docking to drug discovery. Curr Top Med Chem 8, 1555–1572.

Vlachakis, D., Tsagrasoulis, D., Megalooikonomou, V., and Kossida, S. (2013). Introducing Drugster: a comprehensive and fully integrated drug design, lead and structure optimization toolkit. Bioinformatics 29, 126–128.

Warde-Farley, D., Donaldson, S.L., Comes, O., Zuberi, K., Badrawi, R., Chao, P., Franz, M., Grouios, C., Kazi, F., Lopes, C.T., et al. (2010). The GeneMANIA prediction server: biological network integration for gene prioritization and predicting gene function. Nucleic Acids Res 38, W214–220.

Wu, J.J., Zhao, L., Hu, H.G., Li, W.H., and Li, Y.M. (2020). Agonists and inhibitors of the STING pathway: Potential agents for immunotherapy. Medicinal research reviews 40, 1117–1141.

Zhang, C., Hu, Z., Wang, K., Yang, L., Li, Y., Schluter, H., Yang, P., Hong, J., and Yu, H. (2020). Lipidomic profiling of virus infection identifies mediators that resolve herpes simplex virus-induced corneal inflammatory lesions. Analyst 145, 3967–3976.

Zhang, X., Shi, H., Wu, J., Zhang, X., Sun, L., Chen, C., and Chen, Z.J. (2013). Cyclic GMP-AMP containing mixed phosphodiester linkages is an endogenous high-affinity ligand for STING. Molecular cell 51, 226–235.

