## Supplementary figures and images for "Sting orchestrates the crosstalk between polyunsaturated fatty acids metabolism and inflammatory responses"

### Figure S1

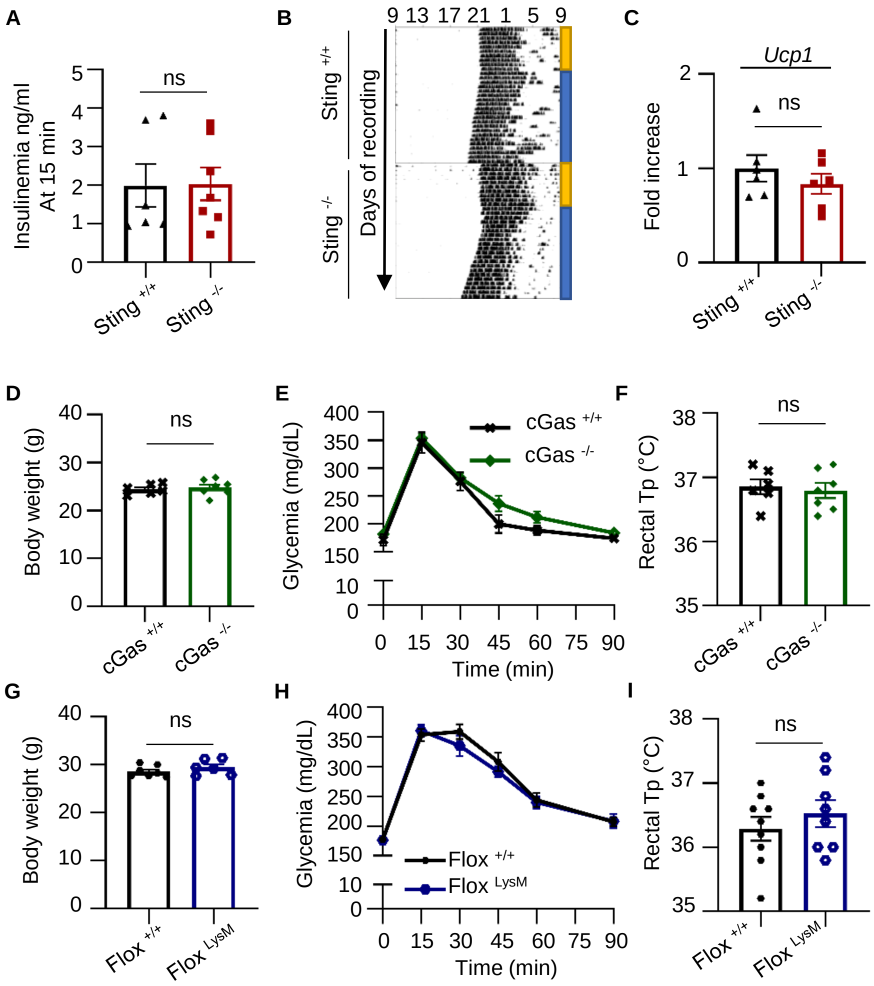

### Figure S2

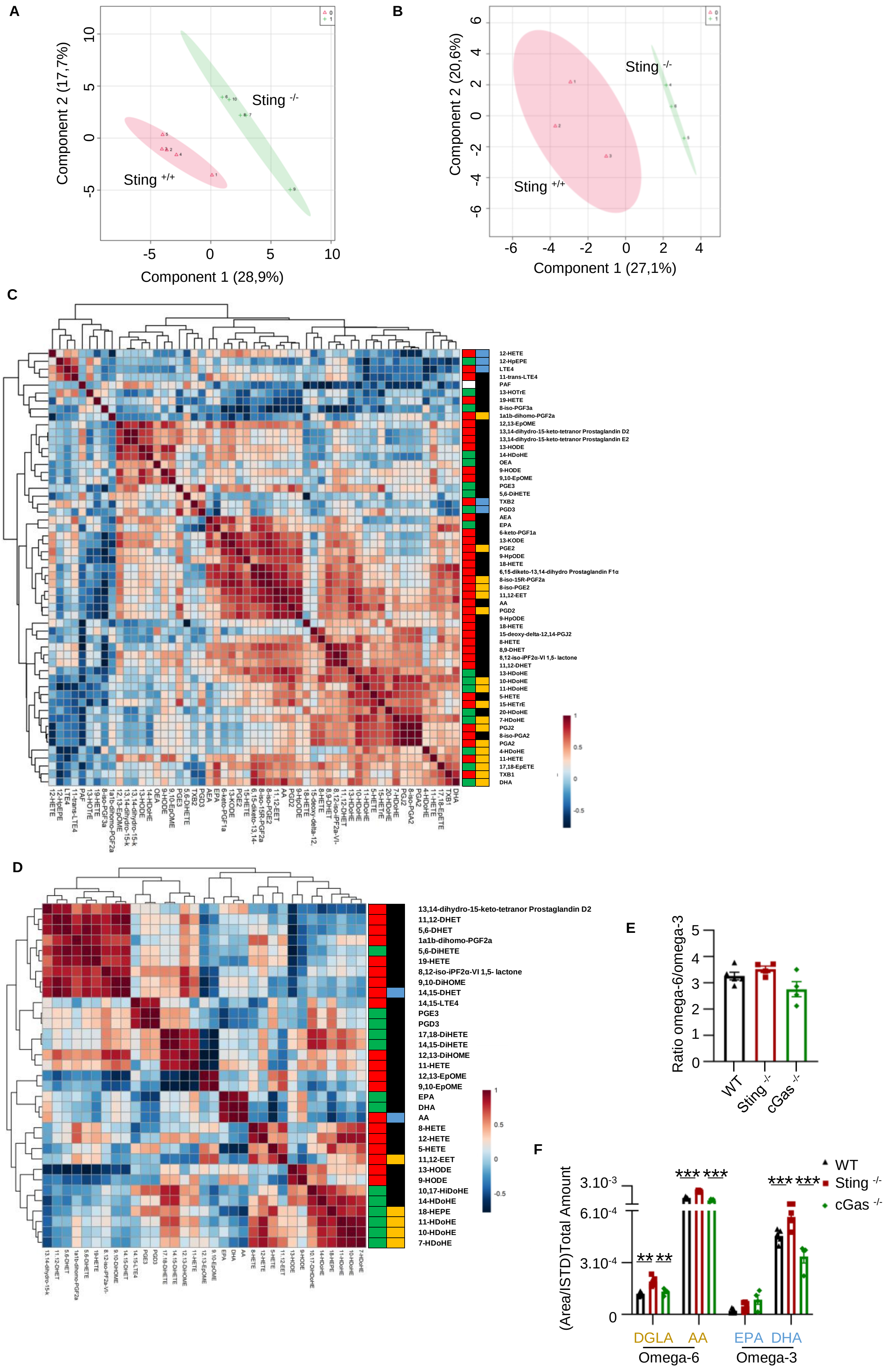

### Figure S3

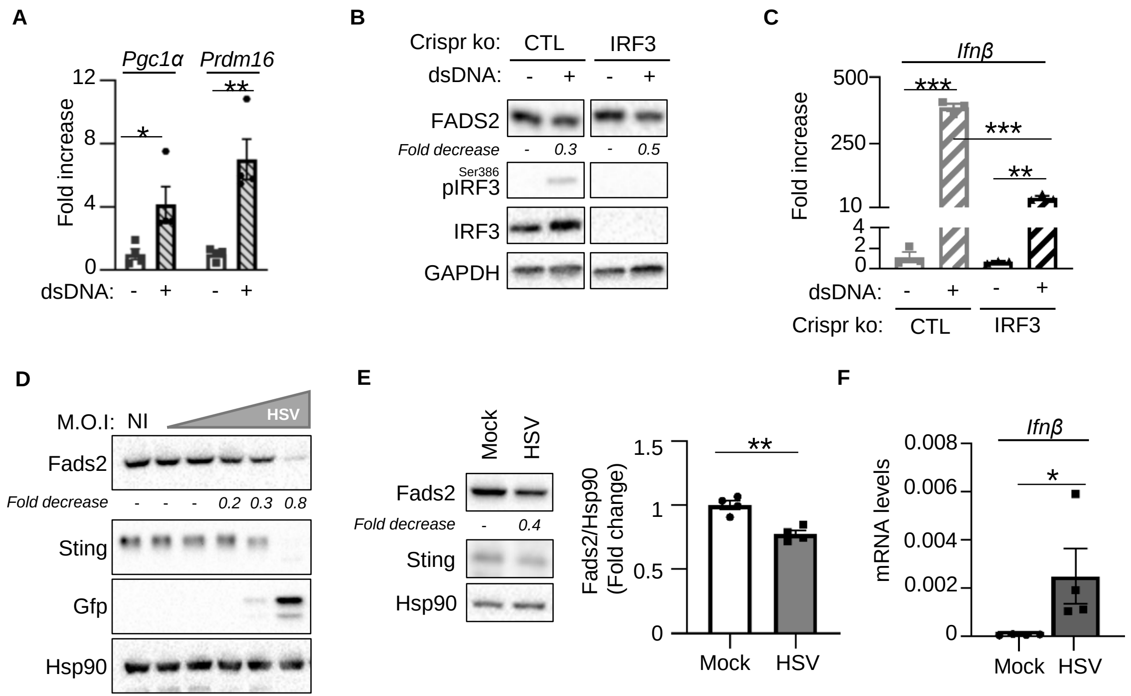

### Figure S4

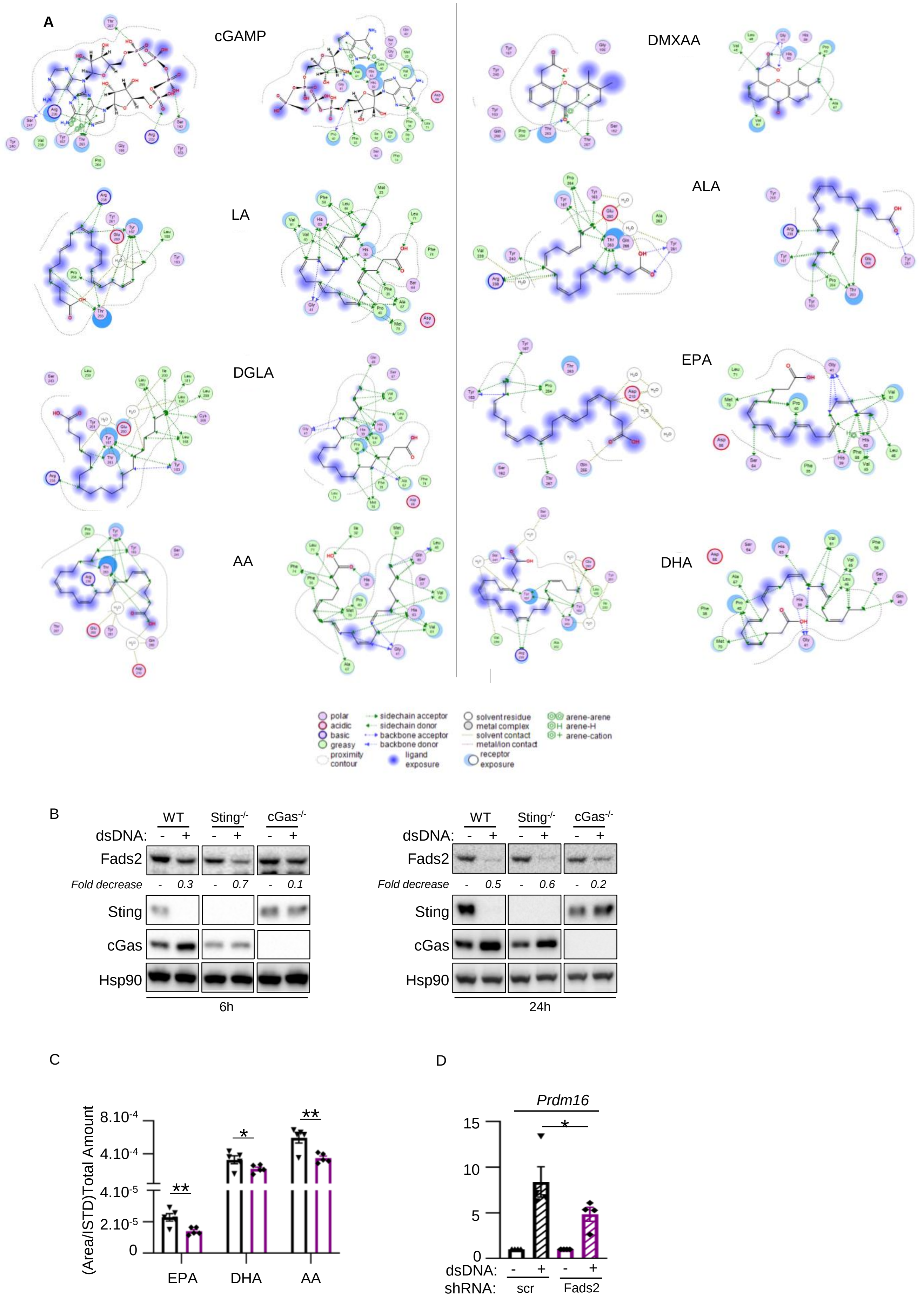
